# The lottery of spermatogenesis

**DOI:** 10.1101/646422

**Authors:** Ata Kalirad, Hadiseh Safdari, Mehdi Sadeghi

## Abstract

The causes of male infertility are yet to be fully understood. Here, we propose that male infertility can be partly attributed to the insufficiency of cytoplasmically-inherited proteins in sperms. Using a simple stochastic model of spermatogenesis, we show that in a range of parameters, the proportion of viable sperms can be reduced simply due to stochastic transmission of proteins from spermatogonia to spermatids.

## Main text

A large proportion of (≈ 50%) infertility in human couples, defined as the inability to achieve pregnancy for 12 months or more [8], can be attributed to men [2]. Since the overwhelming body of work on infertility has been mainly concerned with females, the role of males in infertility is rather unexplored, but there has been an upward trend in studying male infertility in the last decade [9].

A large proportion of (≈ 50%) infertility in human couples, defined as the inability to achieve pregnancy for 12 months or more [8], can be attributed to men [2]. Since the overwhelming body of work on infertility has been mainly concerned with females, the role of males in infertility is rather unexplored, but there has been an upward trend in studying male infertility in the last decade [9].

While a number of factors have been invoked to account for male infertility - from diet to tight underwear (reviewed in [5]) - a far less deterministic explanation might be worth our attention: the stochastic redistribution of spermatogonia proteins among sperms. Can the fickle rules of stochastic sampling result in an sperm with a shortage of proteins, which renders the sperm impotent? While it is generally assumed that 90% of male contribution to infertility is caused by low quality and/or low number of sperms [5], in reality, when taking into the variation within populations, the correlation between sperm quality and time to pregnancy becomes quite murky and the variation in sperm parameters -i.e. sperm concentration, sperm count, etc.- are proved to be quite varied among fertile men [4].

In order to test if a series of cytoplasmic “coin flipping” can result in low quality semen, we simply simulate the distribution of proteins present in a spermatogonia between the sperms that are derived from it. To capture the stochastic distribution of proteins from the differentiated spermatogonia of sperms, we simulate the redistribution of protein 𝒫 in spermatogonia cell in two steps:

1. During meiosis I, *i* number of protein 𝒫 are inherited by the primary spermatocyte *A* such that:

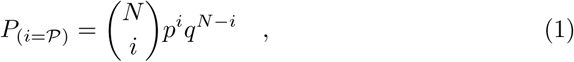

where *N* is the number of protein 𝒫 in the spermatogonia and *p* = *q* = 0.5. After *i* number of 𝒫 are randomly distributed to spermatocyte *A*, spermatocyte *B* inherits *N − i* number of protein 𝒫.
2. Each secondary spermatocyte inherits 𝒫 from its mother cell (i.e., spermatocyte *A* or *B*) according to eq 1, where *N* now is the number of 𝒫 in the mother cell (one of the primary spermatocytes).

The results of a simple model such as this would greatly depend on the number of proteins in the spermatogonia. Probably the most comprehensive measurement of the number of transcripts in differentiated spermatogonia in Human can be found in [3], where single-cell transcript assay was applied to a large number of cells involved in spermatogenesis (Here we are using the mean number of transcripts from 4139 differentiated spermatogonia). Inferring proteome from transcriptome is itself a tad tricky, considering that the estimates for protein/mRNA ranges from 10 to 10^4^ [6]. Here, we have relied on the estimates from an in-depth analysis of protein to mRNA ratios in testis [7]. In addition, we have used the mouse data from [3], as a point of comparison, though in the absence of reliable protein/mRNA data for mice testis, we directly used the number of transcripts as a proxy for the number of proteins (Fig 1).

**Figure 1.**
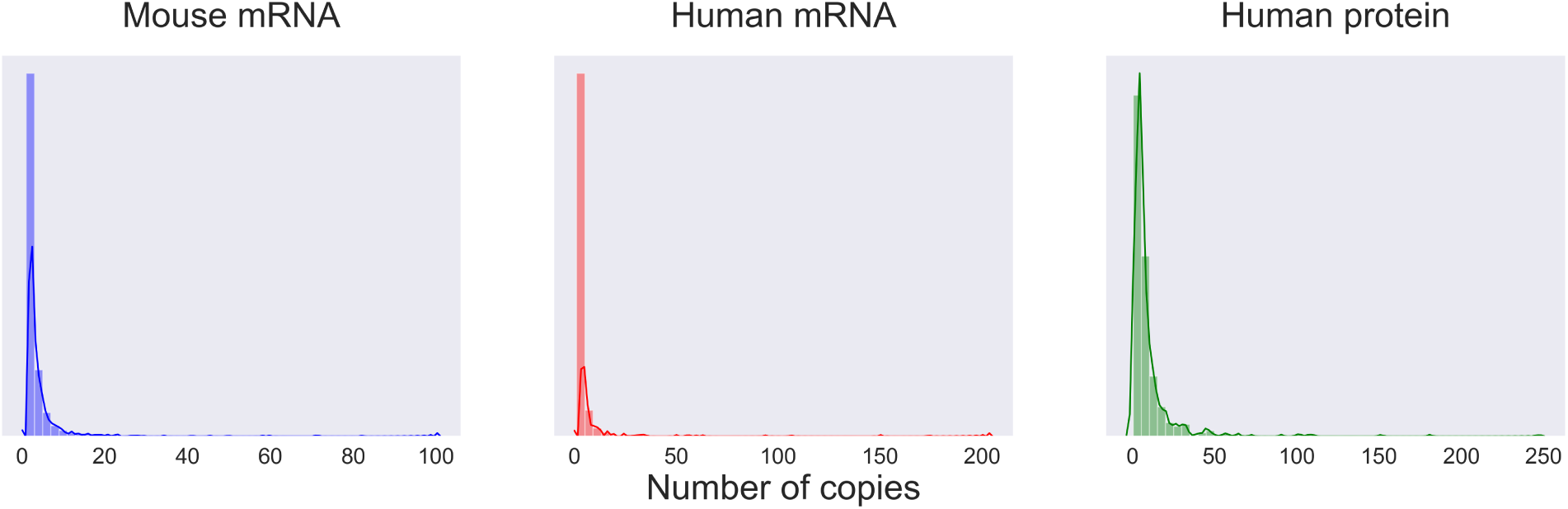
The distribution of the number of copies per transcript and protein in mice and men. The average number of different transcripts in human and mice are based on the single-cell transcriptomics on differentiated spermatogonia (based on data from [3]). To get a better estimate of the number of proteins in human spermatogonia, we combined the transcritpomics data with the protein/mRNA for each gene based on [7].

In our simulations, the number of 𝒫 in each spermatogonia is drawn from a normal distribution with *µ* = *N*_𝒫_, where *N*_𝒫_ is based on the experimental transcriptome and protein/mRNA data, and *σ* = *µ* × 0.2 (0.2 is the reported 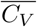 for cell-to-cell protein variability in Human [7]). The viability of sperm *j* is characterized by two parameters: the percentage of protein types required for the functionality of a sperm (𝒩), and the percentage of each functional proteins that has to be inherited from spermatogonia to a sperm to make the sperm viable (𝒯). The proteins included in 𝒩 are randomly chosen from the set of proteins included in this study based on the aforementioned experimental data. The threshold for viability (𝒯) is measured against the reference number of proteins (or transcripts in mice) in the differentiated spermatozoa.

The results from our model indicates the possibility that the sheer accident of not inheriting enough cytoplasmic proteins can lead a large proportion of inviable sperms in man and mice (Fig 2). Understandably, certain parameter combinations -i.e., very high 𝒯 and very high 𝒩 can result in almost no viable sperms while a relaxed combination can produce mostly viable sperms; the reality is almost certainly somewhere in between.

**Figure 2.**
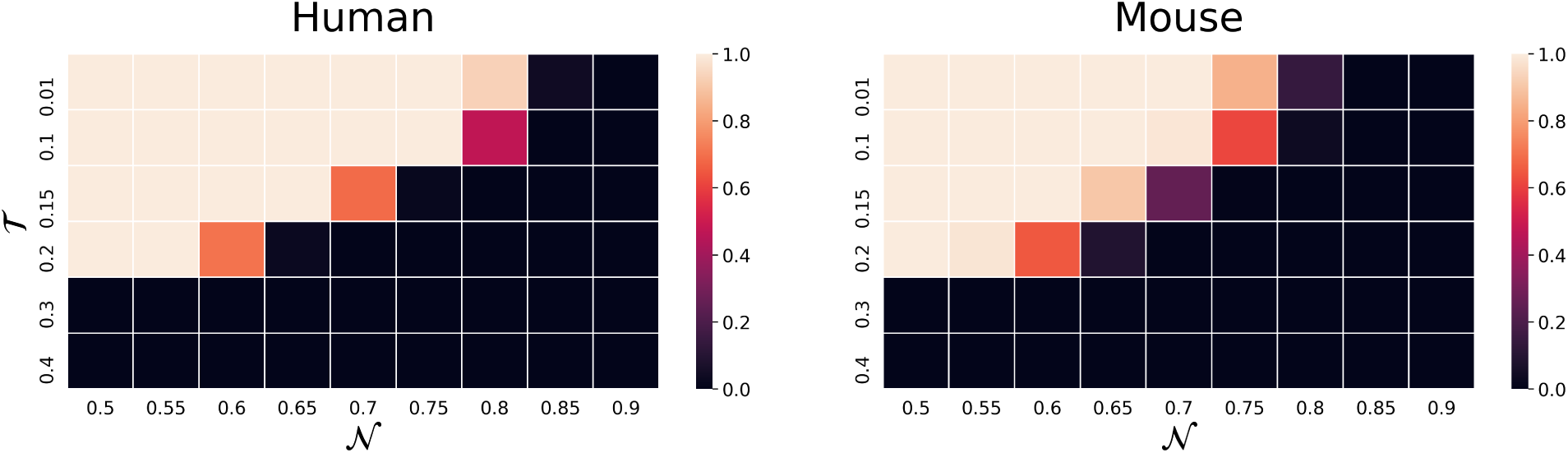
The proportion of viable sperms in man and mice due to mere stochasticity. We measured the proportion of viable sperms by producing 10^5^ sperms with a given 𝒯 and 𝒩. Understandably, very strict combinations of these parameters result in no viable sperm and vice versa.

Our results simply espouse a potential role of stochasticity in the unexplained cases of male infertility but many more questions are yet to be addressed. Our model does not take into account the functionality of proteins nor any clustering between them, two parameters that affect the way proteins distribute between primary and secondary spermatocytes, as well the importance of any given protein for the viability of sperm. Our model is merely a reminder of how mere chance events can lead to biological pattens and can potentially shed some light on the reasons behinds the prevalent suggestion offered by medical specialists that to ensure pregnancy, frequent intercourse (i.e., every day or every other day) is required (e.g., [1]).

## Availability of supporting data

The software supporting the conclusions of this article is available in the https://github.com/Kalirad/Spermatogenesis_Lottery repository.

